# G-quadruplex formation on specific surface-exposed regions of the human ribosomal RNA

**DOI:** 10.1101/435594

**Authors:** Santi Mestre-Fos, Petar I. Penev, Suttipong Suttapitugsakul, Chieri Ito, Anton S. Petrov, Roger M. Wartell, Ronghu Wu, Loren Dean Williams

## Abstract

Profound similarities and critical differences mark ribosomes across phylogeny. The ribosomal core, approximated by the prokaryotic ribosome, is universal, yet mammalian ribosomes are nearly twice as large as those of prokaryotes. Differences in size are due in part to rRNA expansion segments. Here we show rRNA tentacles of Expansion Segment 7 (ES7) of *Homo sapiens* can form G-quadruplexes in vitro. G-quadruplex-forming regions are located on the most surface-exposed regions of the ribosome, near the termini of rRNA tentacles. We characterized rRNA of the large ribosomal subunit by computation, circular dichroism, gel mobility, fluorescent probes, nuclease accessibility, electrophoretic mobility shifts and blotting. We investigated ES7 and oligomers derived from ES7, intact 28S rRNA, and 80S ribosomes and polysomes. We used mass spectrometry to identify proteins that bind to rRNA G-quadruplexes in cell lysates. Proteins that associate with rRNA G-quadruplexes include helicases (DDX3, CNBP, DDX21, DDX17) and heterogeneous nuclear ribonucleoproteins (hnRNPs). And finally, by multiple sequence alignments, we observed that G-quadruplex-forming sequences appear to be a general feature LSU rRNA of the phylum Chordata but not in other phyla. It is known that G-quadruplexes form in telomeres, promoters, and untranslated regions of mRNA but, to our knowledge, they have not been reported previously in ribosomes.

## INTRODUCTION

Cytosolic ribosomes of essentially all extant species contain a ‘common core’ [1] consisting of rRNA and rProteins with universal structure and function. Common core rRNA is reasonably approximated by prokaryotic rRNA; around 90% of prokaryotic rRNA is contained in the common core.

Eukaryotic ribosomes contain additional rRNA and rProteins that form a secondary shell. The rRNA component of the eukaryotic shell contains expansion segments (ESs) that attach to the common core at a handful of specific sites [2-5]. ESs are the most variable rRNA structures over phylogeny, and are known in yeast to play roles in ribosome biogenesis [6] and to bind to chaperone proteins [7, 8]. In chordates, ESs form tentacles (Figure 1) that can extend for hundreds of Ångstroms from the ribosomal surface [9]. The lengths of the tentacles are significantly elongated in mammals and reach a zenith in primates and birds.

Here, we observe the terminal regions of rRNA tentacles of human expansion segments 7 and 27 (ES7_HS_ and ES27_HS_) contain sequences that can form G-quadruplexes. G-quadruplexes are favored by tandem G-tracts separated by short non-specific sequences. ES7_HS_ contains ten tandem G-tracts in one tentacle and four in another. ES27_HS_ contains six tandem G-tracts in *tentacle a*, two groups of three tandem G-tracts in *tentacle b*, and four in helix 63 (Table 1). To investigate G-quadruplex formation in ES7_HS_, we used computation, circular dichroism, gel mobility, fluorescent probing, nuclease accessibility, electrophoretic mobility shift assays (EMSA), dot blotting, Western blotting, and pull-down assays combined with stable isotope labeling with amino acids in cell culture (SILAC) and Mass Spectroscopy. We also performed Multiple Sequence Alignments (MSAs) and database analysis.

Our results indicate that G-quadruplexes can form in a series of model systems including the isolated tandem G-tracts of ES7_HS_, in intact ES7_HS_ and in purified human 28S rRNA. Our data also provide evidence for the formation of G-quadruplexes in 80S ribosomes and polysomes. Pull-down assays show that known G-quadruplex-binding proteins associate with the G-tracts of ES7_HS_. The MSA analyses indicate that G-quadruplex-forming sequences are found in ES7s of all chordate species.

**Table 1.**
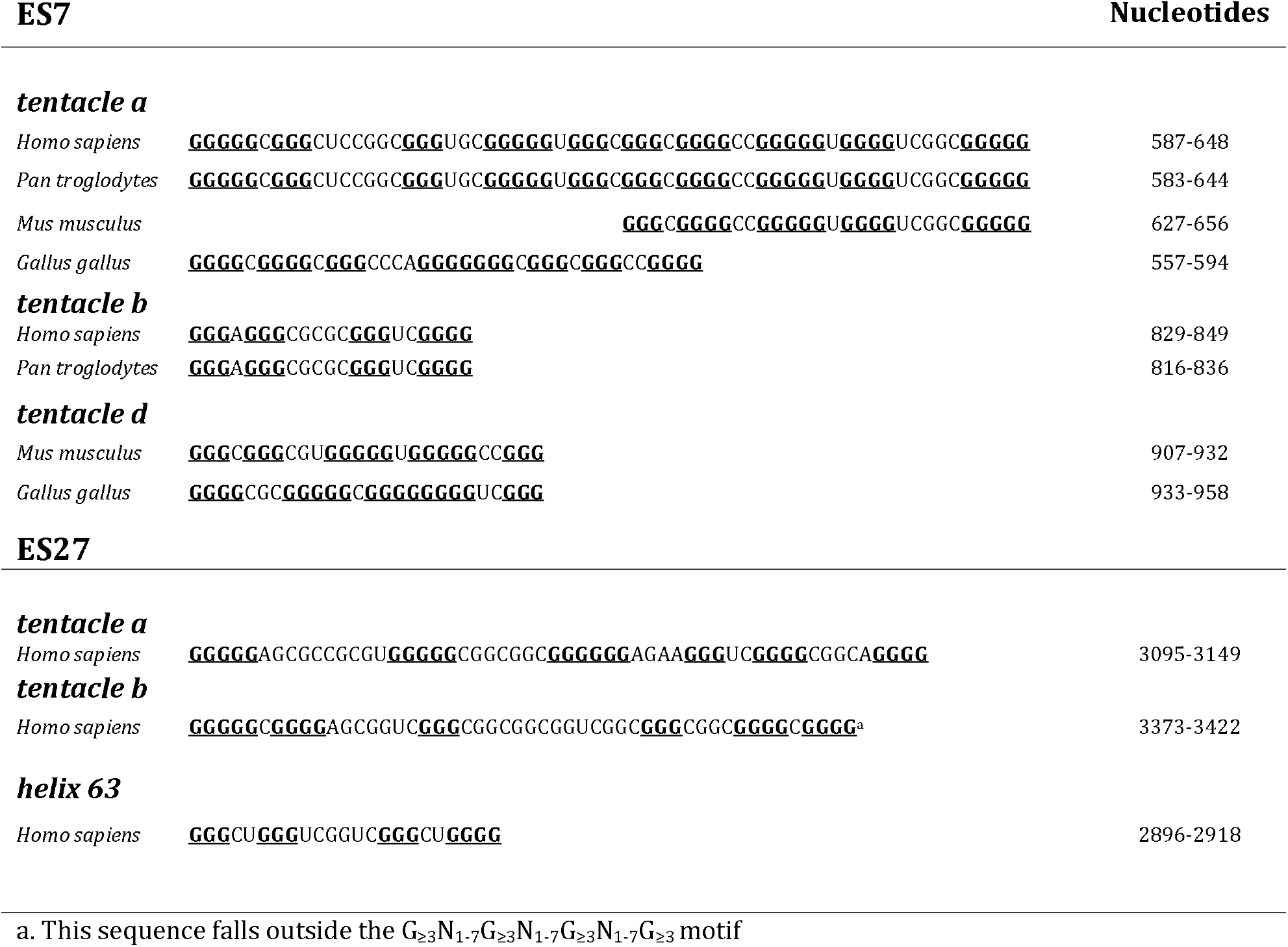
G-quadruplex-forming regions within ES7 and ES27 rRNA

G-quadruplexes have been shown previously to cluster within regulatory RNAs such as 5⍰ and 3⍰ untranslated regions of mRNAs [10] and with the first intron [11]. To our knowledge, this study is the first report of G-quadruplex formation in any ribosomal RNA, which constitutes over 80% of cellular RNA.

**Figure 1.**
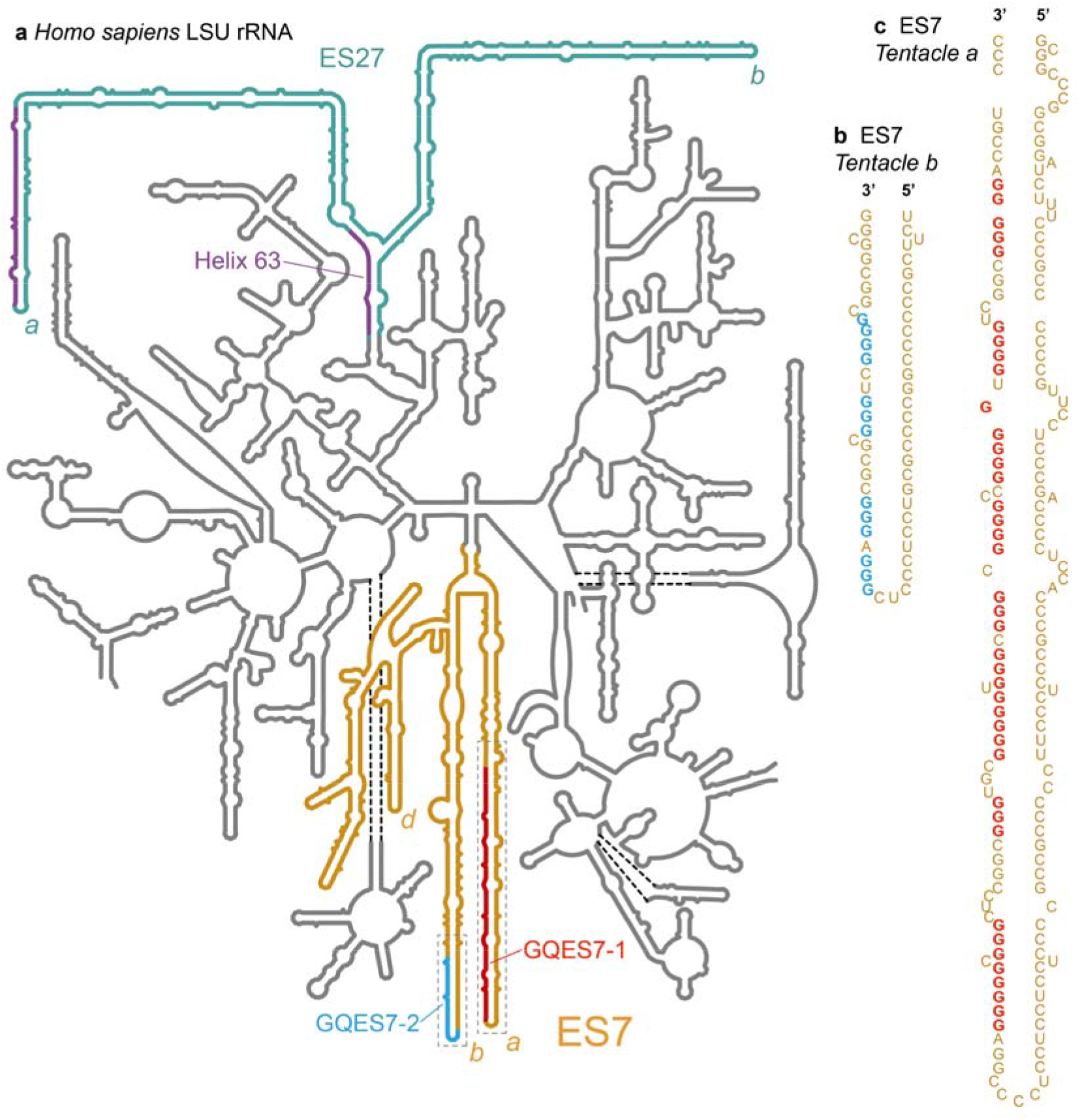
Model secondary structure of the LSU rRNA of *Homo sapiens*. Potential G-quadruplex-forming regions (defined by G_≥3_N_1-7_G_≥3_N_1-7_G_≥3_N_1-7_G_≥3_) are highlighted, a) Expansion segment ES7_HS_ is orange. *Tentacles a, b* and *d* of ES7_HS_ are indicated. G-quadruplex-forming regions GQES7-1 (red) and GQES7-2 (cyan) are highlighted. Expansion segment ES27_HS_ is green with purple G-tracts. Helix 63, at the base of ES27_HS_, contains a G-quadruplex motif (purple). *Tentacles a* and *b* of ES27_HS_ are indicated, b) An expanded view of GQES7-2 indicates the sequence. c) An expanded view of GQES7-1.

## RESULTS

### ES7 and ES27 of the human LSU contain putative G-quadruplex sequences

The propensity of an RNA to form G-quadruplexes can be roughly predicted by numbers and lengths of guanine tracts and the lengths and compositions of loops. The program QGRS Mapper [12] provides “G-scores”, which quantitate this propensity. We have identified ES7 and ES27 as the only human LSU rRNA regions containing sequences reasonably capable of forming these secondary structures. Our computational results suggest that G-quadruplexes can form in the guanine-rich strands at the termini of the longest rRNA tentacles of these two ESs. In tentacles *a* and *b* of ES7_HS_, two regions, here named GQES7-1 and GQES7-2 (Figure 1), meet the G-quadruplex consensus (G_≥3_N_1-7_G_≥3_N_1-7_G_≥3_N_1-7_G_≥3_). The G-scores of GQES7-1 and GQES7-2 are in the range of well-established RNA G-quadruplexes.

The sequence 5’ <u>GGGG</u>CC<u>GGGG</u>GU<u>GGGG</u>UCGGC<u>GGGG</u> 3’ (nts 623-647, within GQES7-1, Figure 1, Table 1) gives a G-score of 60. The sequence 5’ <u>GGG</u>UGC<u>GGG</u>GGU<u>GGG</u>C<u>GGG</u> 3’ (nts 603-621, also within GQES7-1) gives a G-score of 40. The sequence 5’ <u>GGG</u>A<u>GGG</u>CGCGC<u>GGG</u>UCG<u>GGG</u> 3’ (nts 829-849 within GQES7-2, Figure 1, Table 1) gives a G-score of 38. The difference between the G-scores of GQES7-1 and GQES7-2 suggests more stable and/or more extensive G-quadruplex formation in GQES7-1 than in GQES7-2. A greater propensity of GQES7-1 over GQES7-2 for G-quadruplex formation is seen in all experiments below.

As a positive control for our computations and experiments, we used the ADAM10 G-quadruplex. This stable and well-characterized G-quadruplex-forming RNA, found in the 5’-UTR of the mRNA of ADAM10 metalloprotease [13], gives a G-score of 42. As negative controls, we used two mutant RNA oligomers (*mt*ES7-1 and *mt*ES7-2) that are analogous to GQES7-1 and GQES7-2 in composition and length, with disrupted G-tracts (Table S.l). Neither gives a G-score.

Near the terminus of human ES27 (ES27_HS_, Figure 1, Table 1), the sequence 5’ <u>GGG</u>GAGAA<u>GGG</u>UCG<u>GGG</u>CGGCA<u>GGG</u> 3’ (nts 3124-3148, *tentacle a*) gives a G-score of 40. ES27_HS_ also contains a putative G-quadruplex-forming region in Helix 63 near the junction of two tentacles. In this study, we focused only ES7_HS_. However, based on the high correlation of G-scores and our experimental observation of G-quadruplexes within ES7_HS_, we expect G-quadruplexes to also form in ES27^HS^.

We investigated the ability of the two regions within ES7_HS_ (GQES7-1 and GQES7-2, Figure 1) to form G-quadruplexes *in vitro* by several methods.

### ThT fluorescence

ThT is known to yield intense fluorescence at 487 nm upon association with G-quadruplexes [14, 15]. ThT results here suggest formation of G-quadruplexes in GQES7-1, GQES7-2 and intact ES7_HS_ in the presence of K^+^, the monovalent ion thought to stabilize G-quadruplexes (Figure 2a). The signal for GQES7-2 is attenuated compared to that of GQES7-1. This difference, which suggests less stable G-quadruplexes and/or less extensive G-quadruplex formation of GQES7-2, is consistent with the computed propensity and is observed here in a variety of assays. The formation of G-quadruplexes by GQES7-1, GQES7-2 and intact ES7_HS_ is supported by competition assays with pyridostatin (PDS) (Figure 2c and 2d). PDS is a G-quadruplex stabilizer and a ThT competitor with a greater affinity than ThT for G-quadruplexes [16]. In this series of experiments ADAM10 was used as a positive control and *mt*ES7-1, *mt*ES7-2 and tRNA were used as negative controls.

### Circular dichroism

CD has been used widely to study RNA and DNA G-quadruplexes. Our CD spectra of GQES7-1 and GQES7-2 are consistent with formation of parallel G-quadruplexes based on a characteristic peak at 260 nm and a trough at 240 nm [17-21] (Figure 2b). G-quadruplexes are thought to be more stable in K^+^ than in Li^+^ or Na^+^ [22]. The intensity of the 260 nm peak of GQES7-1 is attenuated by about 15% when the monovalent cation is switched from K^+^ to Na^+^ or Li^+^ (not shown).

**Figure 2.**
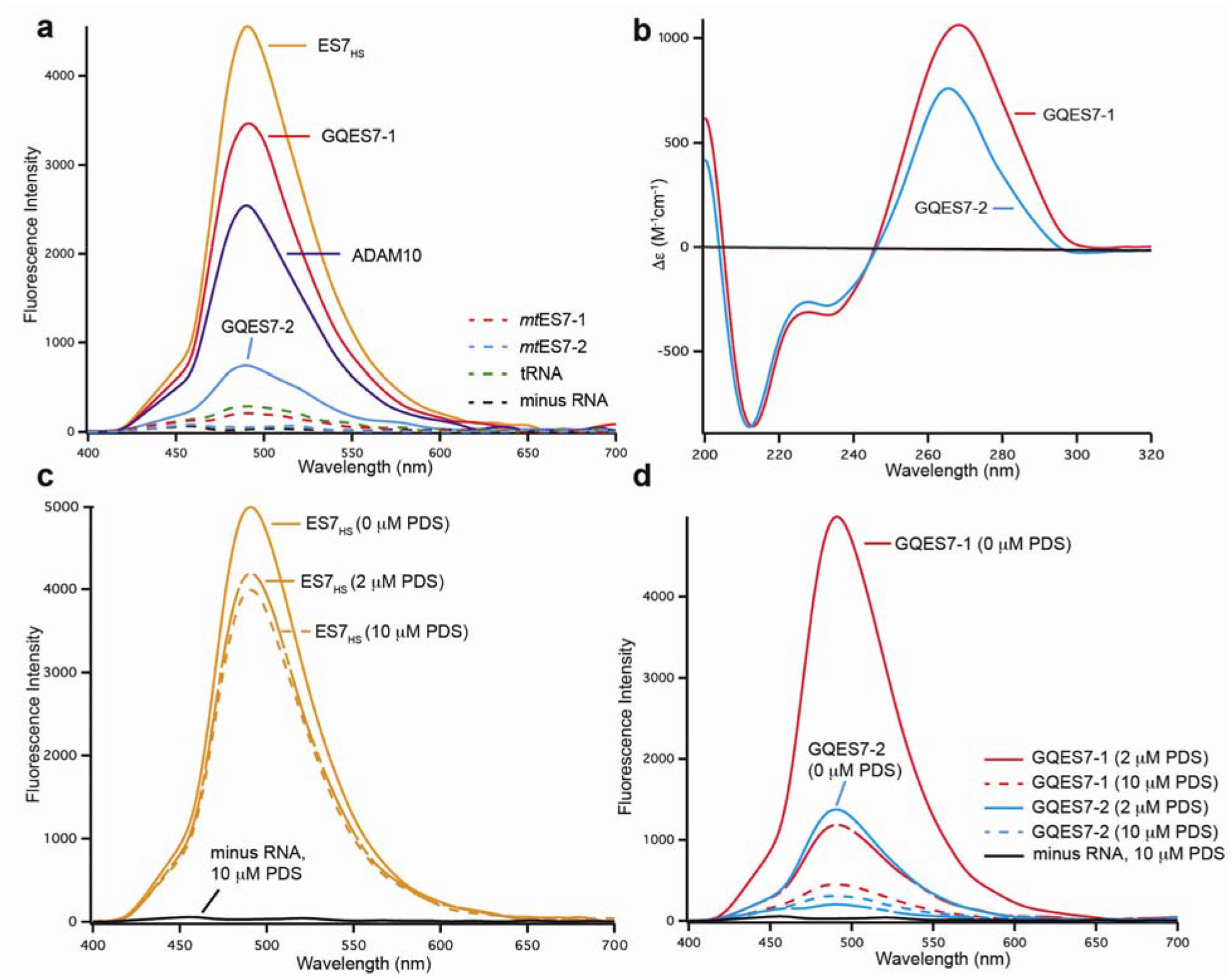
Formation of G-quadruplexes in GQES7-1, GQES7-2, and ES7_HS_. a) Fluorescence emission of ThT in the presence of ES7_HS_, GQES7-1, GQES7-2, or ADAM10. Negative controls (dashed) are tRNA, *mt*ES7-1, *mt*ES7-2 and minus RNA. b) CD spectra of GQES7-1 and GQES7-2. In the presence of ThT, PDS was added to c) intact ES7_HS_ and to d) GQES7-1 and GQES7-2.

### Gel staining

ThT staining of GQES7-1 and GQES7-2 after native gel electrophoresis in the presence of Na^+^ or Li^+^ or K^+^ is in agreement with the CD data (Figure 3a and 3b). The RNAs are expected to fluoresce within the gel only upon formation of G-quadruplexes. GQES7-1 and GQES7-2 fluoresce more intensely in the presence of K^+^ than in Na^+^ or Li^+^. Consistent with computation, ThT fluorescence in solution, and CD spectroscopy, GQES7-1 forms more extensive G-quadruplexes than GQES7-2 in the gel staining experiments.

**Figure 3.**
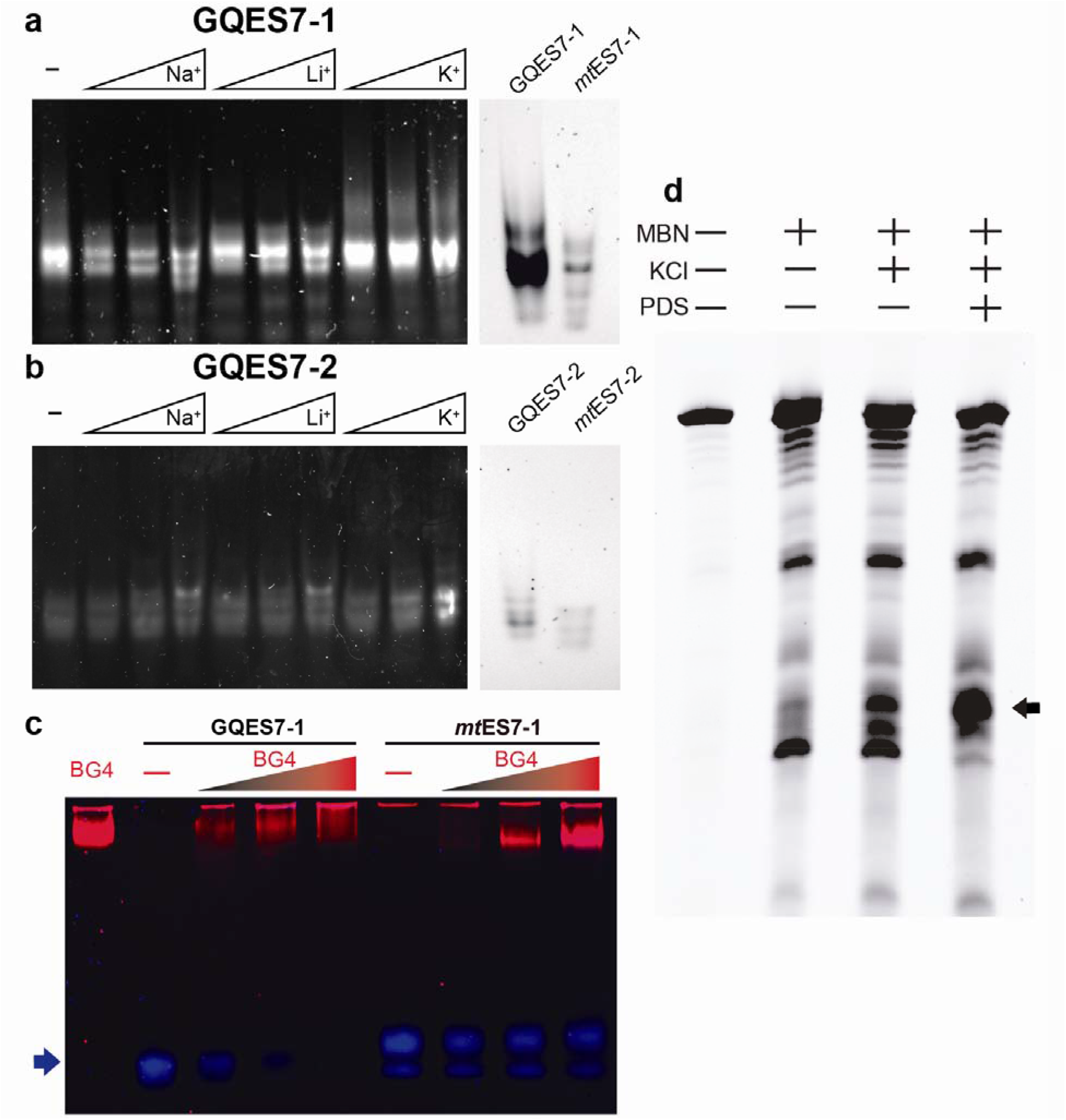
The effect of monovalent ion type on G-quadruplex formation. Shown here are a) GQES7-1 and b) GQES7-2 RNAs with varying concentrations of Na^+^, Li^+^, and K^+^ (50,100, 250 mM), resolved on a native gel stained with ThT. The mutants *mt*ES7-1 and *mt*ES7-2 do not fluoresce when stained with ThT, indicating the lack of G-quadruplex formation in these RNAs (gels on the right), c) EMSA of the BG4 antibody with GQES7-1 and its mutant *mt*ES7-1, visualized on a native gel. GQES7-1 and *mt*ES7-1 RNAs were loaded at a constant strand concentration with increasing concentrations of BG4 antibody. The RNA (arrow) is blue and the protein is red. d) ES7_HS_ cleavage by mung bean nuclease. ES7 was annealed with or without KCl and with or without PDS. The black arrow indicates cleaved rRNA.

### Mung bean nuclease cleavage

The regions of 28S rRNA capable of forming G-quadruplexes are opposed by complementary C-rich strands (Figure 1). Upon G-quadruplex formation from duplex rRNA, the C-rich strand would be forced to dissociate to a single strand. To confirm this model, we examined cleavage of ES7_HS_ by mung bean nuclease (MBN) (Figure 3d), which preferentially cleaves single-stranded RNA and DNA. The results here show that MBN cleaves ES7_HS_ under G-quadruplex stabilizing conditions to a greater extent than under conditions that destabilize G-quadruplexes (Figure 3d). Addition of K^+^ increases the extent of cleavage and addition of PDS to K^+^ further increases the extent of cleavage. The simplest interpretation of these results is that ES7_HS_ is dynamic and can exist as a mixture of duplex and G-quadruplex forms. As a negative control, extent of MBN hydrolysis of tRNA did not increase upon addition of K^+^ and/or PDS (Figure S.2).

### Antibody binding

BG4 is an antibody developed by Balasubramanian and coworkers [23, 24] that binds to a variety of G-quadruplex types but not to other nucleic acids such as RNA hairpins, single-stranded or double-stranded DNA. Here, an EMSA was performed with BG4 to test for G-quadruplex formation in GQES7-1 (Figure 3c). BG4 was also used for dot blotting experiments with GQES7-1, GQES7-2, and intact ES7_HS_ (Figure 4). We observe binding of GQES7-1, GQES7-2, and intact ES7_HS_ with BG4. Consistent with the results above, GQES7-1 binds more tightly than GQES7-2 to BG4, which is close to the negative controls (Figure 4a and 4c).

The experiments presented above confirmed that the sequences within human rRNA ES7 of the 28S rRNA are capable of forming G-quadruplexes. We have investigated whether the 28S rRNA obtained from cells could form G-quadruplexes when this RNA is rProtein-free or assembled in ribosomes.

The 28S rRNA was extracted from HEK293T cells and dot blotting was performed with BG4 (Figure 4d). The results indicate that 28S rRNA forms G-quadruplexes *in vitro*. To confirm that the 28S rRNA is capable of forming G-quadruplexes when assembled in the intact ribosome, dot blotting was also performed with purified 80S human ribosomes and polysomes (Figure 4e and 4f).

The results show that the BG4 antibody selectively binds to intact human ribosomes and polysomes in a concentration-dependent manner. PDS enhances binding of the antibody, as expected. The more extensive binding of the antibody to polysomes than to monomer ribosomes suggests the possibility of formation intermolecular of G-quadruplexes in polysomes.

**Figure 4.**
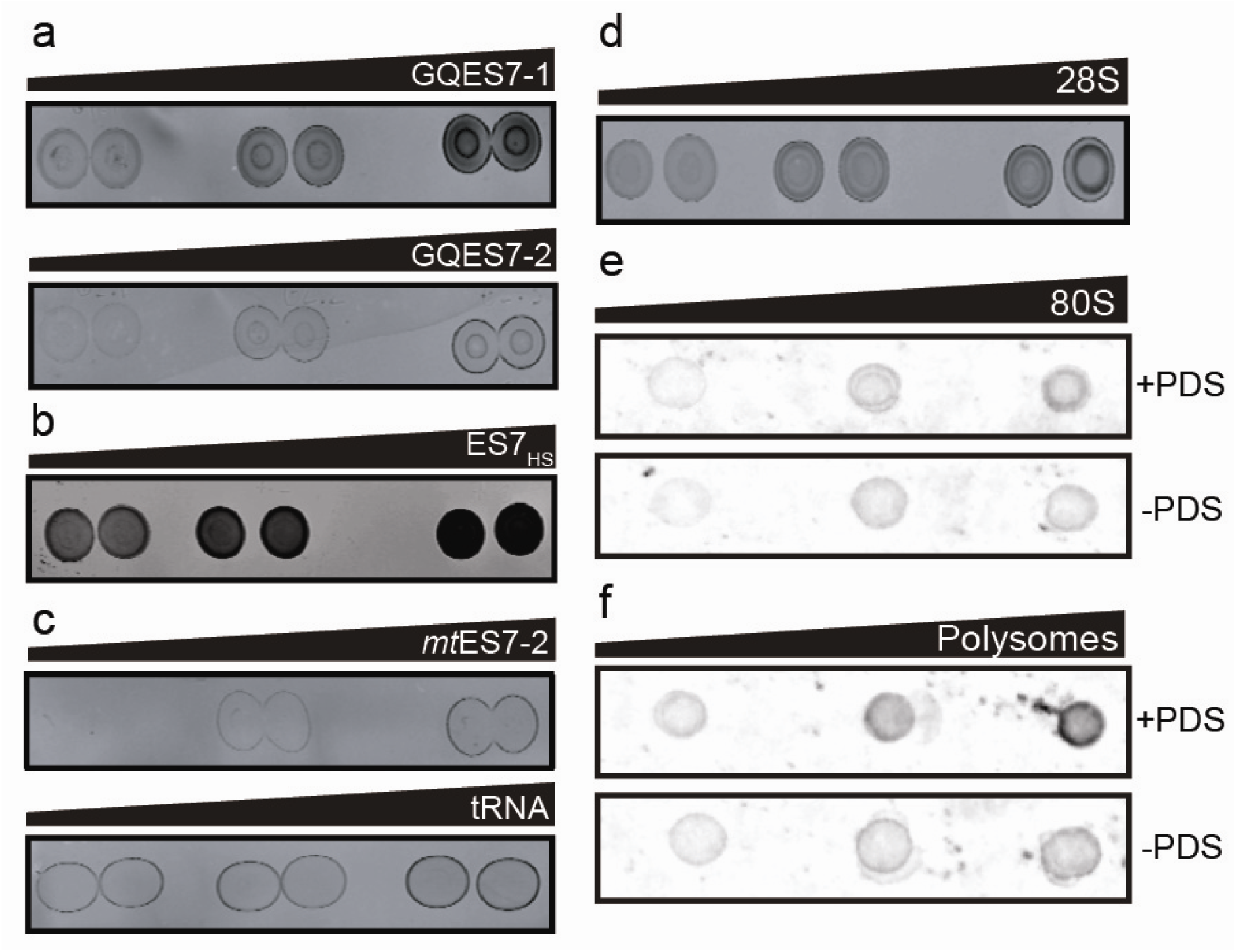
Dot blots performed with the BG4 antibody on a) GQES7-1 and GQES7-2, b) intact ES7_HS_, c) the negative controls *mt*ES7-2 and tRNA, d) the 28S rRNA extracted from HEK293T cells and on e) human 80S ribosomes and f) polysomes purified from HEK293 cells. All samples were incubated in the presence of 50mM KCl and ribosomes and polysomes were further analyzed with or without 10 μM PDS, which stabilizes G-quadruplexes. Samples were loaded onto the membrane in increasing amounts from left to right.

### rRNA G-quadruplex-forming sequences are observed throughout the Chordata phylum

Focusing specifically on translation we have developed the SEREB Database [1], which contains fully curated sequences of rRNAs from all major phyla, yet samples the tree of life in a sparse, efficient and unbiased manner. Here we extended the SEREB database from 10 chordate species to 17 for a fine-grained analysis of ES7.

#### G-quadruplex-forming sequences in chordate ES7s

Our MSA confirms that the lengths of rRNA tentacles of eukaryotes are variable, reaching maxima in species such as *G. gallus* and *H. sapiens*. Sequences of the relevant segments of ES7s of various eukaryotes are illustrated in Figure 5, demonstrating the conservation of G-quadruplex-forming sequences motif [G_≥3_N_1-7_G_≥3_N_1-7_G_≥3_N_1-7_G_≥3_ (number of G tracts (n) >3)] and their preferential locations near the termini of tentacles. The number of tandem G-tracts in *tentacle a* of ES7 is ten in human and chimpanzee and eight in rat and chicken. Fish reptiles and amphibians appear to lack the G-tract motif described here. However, G-tracts outside of the motif can form G-quadruplexes [25, 26]. It is possible that G-tracts with n<4 form intermolecular G-quadruplexes with other tentacles or with other ribosomes as in polysomes.

#### G-quadruplex-forming sequences in chordate ESs other than ES7

Our analysis of the extended SEREB Sequence Database suggests that G-quadruplex-forming sequences are universal to chordates (Table S.2). Several chordate species present G-quadruplex-forming sequences in tentacles other than ES7 (Table S.2).

**Figure 5.**
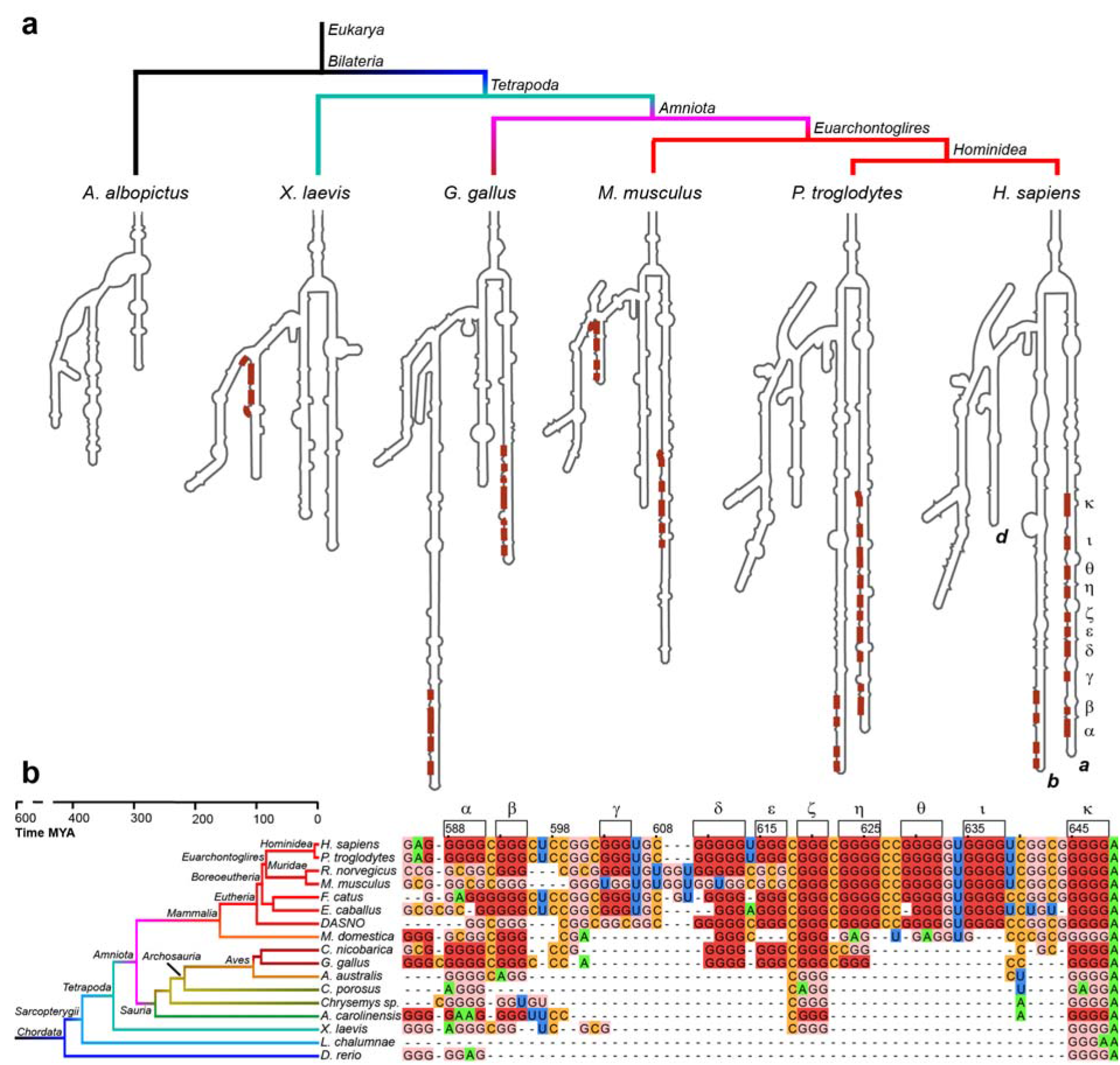
G-tracts in *tentacle a* are observed in birds and mammals. a) Secondary structural models of ES7 from various eukaryotes. G-tracts with G_≥3_N_1-7_G_≥3_N_1-7_G_≥3_N_1-7_G_≥3_ are highlighted in red. b) Sequence alignment of *tentacle a* showing conservation of G-quadruplex-forming sequences in chordates. G-tracts in both panels are labeled with Greek symbols. Nucleotides are colored by type. G’s within G-tracts are dark red. Other G’s are light red. All nucleotides are numbered in accordance with *H. sapiens* 28S rRNA. Sizes of eukaryotic ES7 secondary structures are not to scale. Complete species nomenclature is provided in Table S.3.

#### Absence of G-quadruplex sequences in non-chordate rRNAs

To determine the phylogenic distribution of G-quadruplexes in LSU rRNA, we inspected highly curated sequences of 20 non-chordate eukaryotes from the SEREB database [1]. Thus far we can find no evidence of G-quadruplex-forming sequences in non-chordate eukaryotes.

### RNA remodeling proteins bind selectively to rRNA G-quadruplexes

The localization of rRNA G-quadruplex sequences on ribosomal surfaces would suggest they interact with non-ribosomal proteins. To identify the proteins that bind to rRNA G-quadruplexes, we performed pull-down experiments using stable isotope labeling with amino acids in cell culture (SILAC). We focused on GQES7-1, the longest and most stable G-quadruplex-forming region in human rRNA (Figure 6). GQES7-1 rRNA was linked on the 3’ end to biotin (GQES7-1-Biotin) and interacting proteins from human cell lysates were pulled down and analyzed by mass spectrometry. The biotinylation of GQES7-1 did not disrupt the G-quadruplexes (Figure S.3). Several known G-quadruplex-binding proteins were pulled down by this assay (CNBP, YBOX1, hnRP F, hnRP H, DDX21, DDX17, DDX3X) [27-33]. A significant number of helicases were identified (DDX3, CNBP, DDX21, DDX17), all of which have been reported to unfold G-quadruplexes. In addition, a significant number of heterogeneous nuclear ribonucleoproteins (hnRNPs) were bound to GQES7-1, including hnRNP G-T/RMXL2, hnRNP M, hnRNP G/RBMX, hnRNP H2, hnRNP H, hnRNP F, hnRNP H3, and FUS. hnRNPs are a family of RNA-binding proteins with functions including pre-mRNA processing and transport of mRNAs to ribosomes [34]. Several of these proteins have been previously identified as ribosome-binding proteins [35].

To support results of the pull-down experiments, Western blotting was performed with four of the resulting proteins (Figure 6d). We assayed a DEAD-box RNA helicase (DDX3X), a heterogeneous nuclear ribonucleoprotein (hnRNP H), the RNA-binding protein FUS, and a pre-mRNA polyadenylation stimulator (FIP1). hnRNP H and DDX3X have been previously identified as G-quadruplex-binding proteins. However, all four proteins are seen to bind to GQES7-1 in the Western blot, suggesting we have tapped an uncharacterized pool of G-quadruplex-binding proteins.

**Figure 6.**
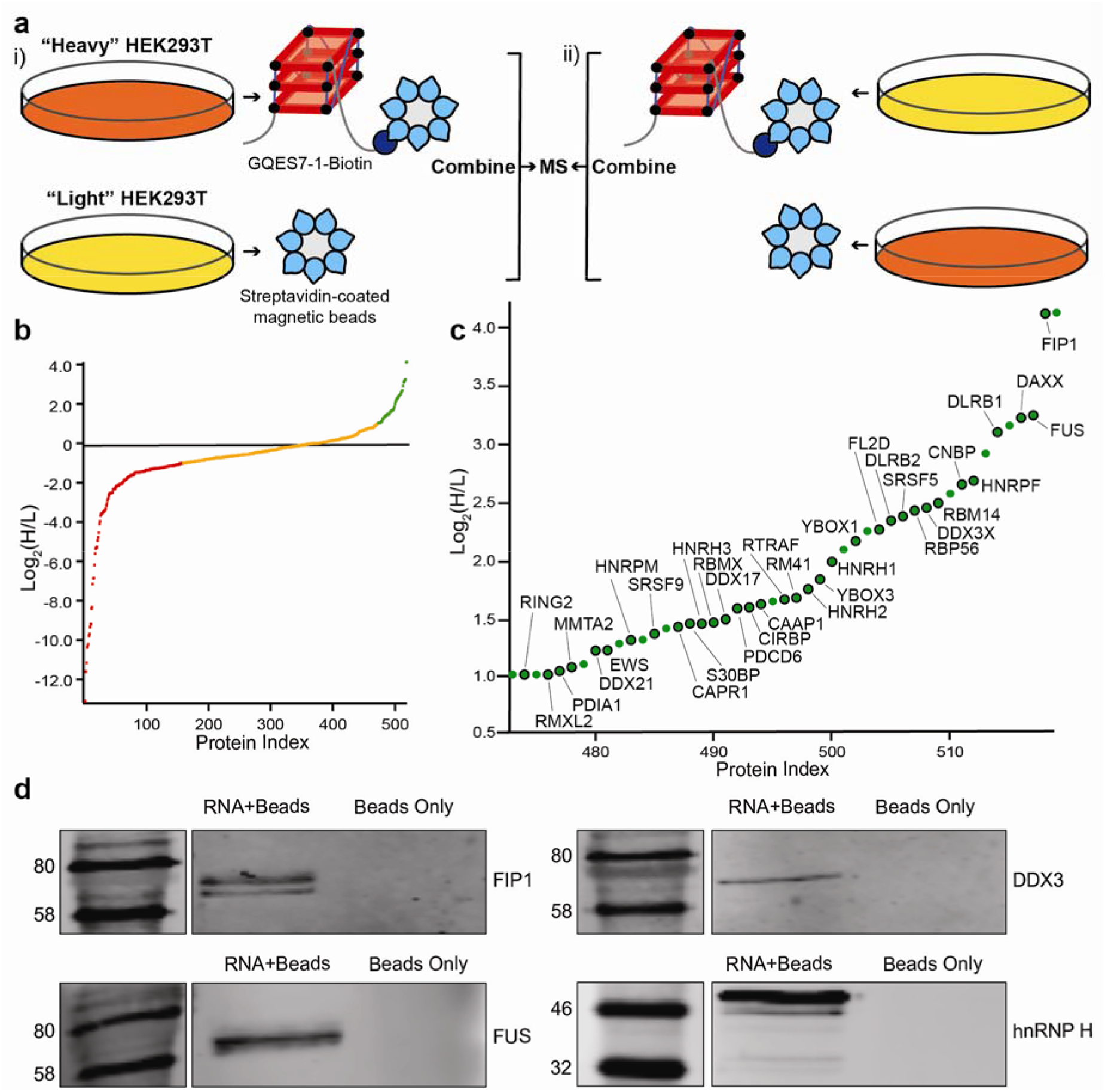
Identification of GQES7-1-binding proteins. a) Scheme of SILAC experiment. “RNA+beads” samples were combined in HEK293T grown in heavy media (i). The “Beads Only” control sample was combined in HEK293T grown in light media. To verify the proteins identified by this method, the experiment was performed using reverse labeling (ii). b) Scatter plot representing fold enrichment of the proteins binding to GQES7-1 in “Heavy” HEK293T. Color representation indicates specific proteins that bound more tightly to GQES7-1 than to the beads (green), to the beads than to GQES7-1 (red) or bound to the beads and GQES7-1 to a similar extent (orange). c) A close-up of the green region of the scatter plot represented in b). Dots with a black contour are used to indicate proteins that appeared in the green region of the two replicate experiments described in a). d) Western blotting analyses of the eluted proteins from the RNA pull-down of HEK293T. All four blotted proteins (FIP1, FUS, DDX3 and hnRNP H) eluted from the GQES7-1 sample (RNA+Beads) but not from the control (Beads Only), confirming the SILAC results.

## DISCUSSION

RNA G-quadruplexes have been studied predominantly in the context of mRNAs, where they have been shown to help regulate expression of certain proteins. The results presented in this study indicate that the human LSU rRNA, one of the most abundant RNAs, contains sequences capable of forming G-quadruplexes (Figure 7). Computation, *in vitro* ThT fluorescence, CD spectroscopy, EMSAs, nuclease digestion and blotting with a G-quadruplex antibody provide a consistent picture of the propensities of G-quadruplex formation in the human 28S rRNA. Our phylogenetic analysis suggests that the 28S rRNAs of all chordate ribosomes contain G-quadruplex-forming sequences near the termini of ES7 *tentacle a*. Our results suggest that nature’s most complex organisms have evolved long rRNA tentacles with unexpected polymorphism, including the ability to convert to G-quadruplexes. At the limit, these sequences can be hundreds of Ångströms from the ribosomal core, suggesting roles in recruiting specific proteins or in stabilizing polysomes.

G-quadruplex-forming rRNA sequences appear to be a general feature of ribosomes of the phylum Chordata. We have inferred this by multiple sequence alignments. The specific sequences and exact locations of the G-quadruplexes on the tentacles are variable across phylogeny. We searched the SEREB database and could find no evidence of G-quadruplex-forming sequences outside of the Chordata phylum. The SEREB database is specifically designed for rRNA analysis, and includes species from all major phyla, and samples the tree of life in a sparse, efficient and accurate manner [1]. It contains complete and highly curated rRNA sequences.

To our knowledge, the possibility that rRNA can be loci of G-quadruplexes has not been investigated previously. G-quadruplex-forming sequences have been described in genes encoding for rRNA, where they are proposed to influence transcription [36] and bind to the nucleolar protein nucleophosmin [37]. However, these studies have focused on the external and internal transcribed regions (ETS and ITS) and are not part of the assembled ribosome. RNA G-quadruplexes have been observed previously in the cytoplasm of human cells [23] and are known to bind tightly to a wide variety of proteins.

**Figure 7.**
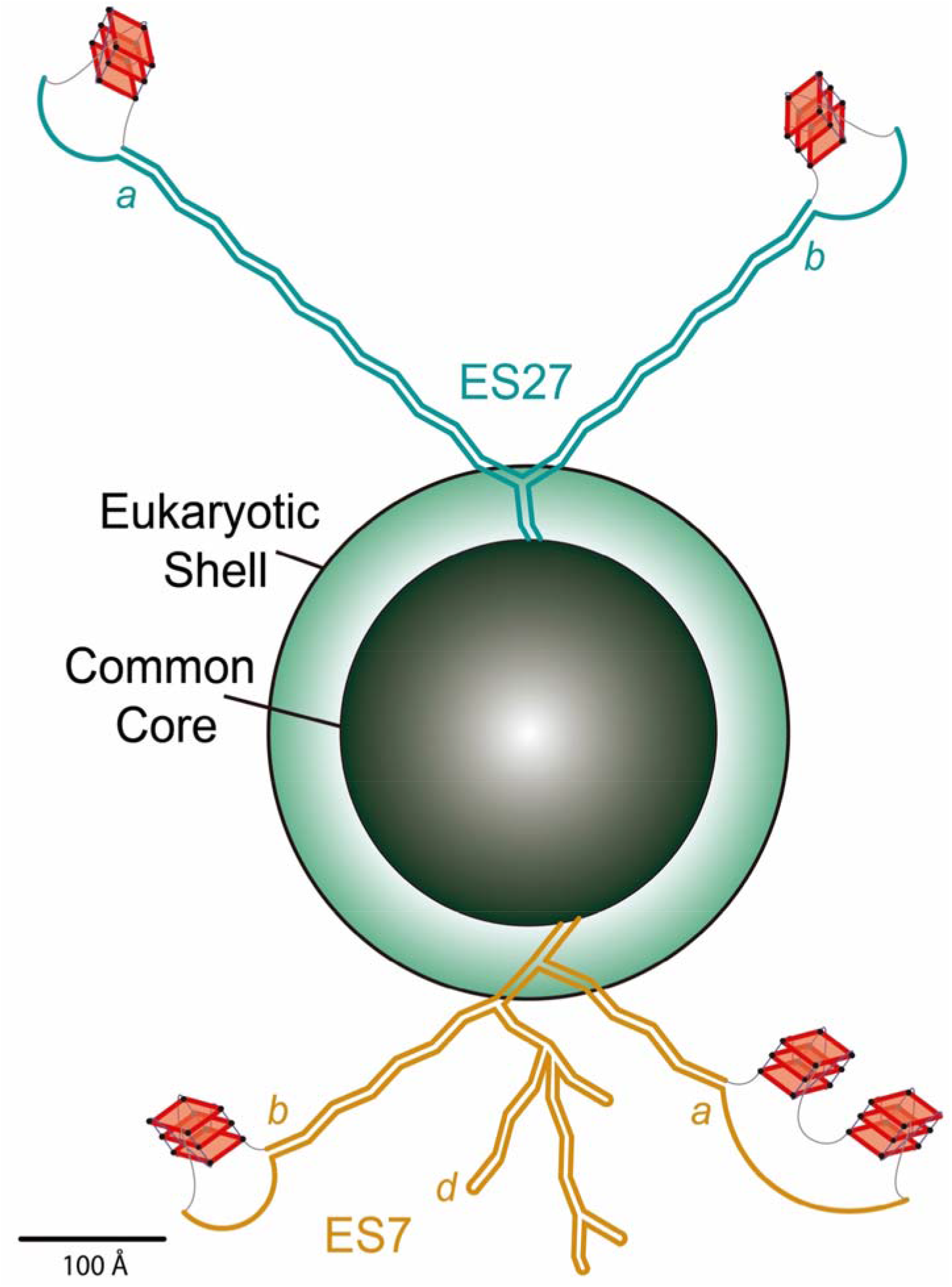
Schematic representation of the common core, the eukaryotic shell and the tentacles of metazoan ribosomes. G-quadruplexes are indicated on the tentacles of ES7 and ES27 in the LSU of the *Homo sapiens* ribosome. The lengths of ES7_HS_ (orange) and ES27_HS_ (green) tentacles are roughly scaled to the size of the common core. *Tentacles a, b*, and *d* (ES7_HS_) and *tentacles a* and *b* (ES27_HS_) are indicated. *Tentacle a* of ES7 is represented as two G-quadruplexes based on our results, which indicate more extensive and/or more stable G-quadruplexes than those found in *tentacle b*. The G-quadruplex region found in Helix 63 of ES27_HS_ is not represented here. The G-quadruplexes represented in *tentacle b* of ES27_HS_ do not fall within the G_≥3_N_1-7_G_≥3_N_1-7_G_≥3_N_1-7_G_≥3_ motif but this region contains extensive G-tracts (Table 1) that could form G-quadruplexes.

The extent to which G-quadruplexes form in the context of cellular environments remains uncertain. Bartel and coworkers have suggested that mRNA G-quadruplexes are globally unfolded in eukaryotic cells, presumably by a mechanism involving regulated unwinding factors [38]. These experiments focused on poly-adenylated mRNAs rather than rRNAs, but these unwinding factors may influence G-quadruplexes in rRNA as well.

The preferential localization of rRNA G-quadruplex regions near the termini of specific tentacles of ES7 and ES27 suggests these regions as loci for specific cytosolic proteins. Here we identified multiple human RNA helicases and other RNA remodeling proteins that bind to rRNA G-quadruplexes. These proteins could be participants in G-quadruplex regulation on ribosomes.

Our observation that polysomes appear to form more extensive G-quadruplexes than monomer ribosomes suggests a role for intermolecular G-quadruplexes in closely associated ribosomes. Our work points to the possibility that, inside cells, ribosomes present polymorphic tentacles that can switch between unimolecular and multimolecular G-quadruplexes and duplex forms. In this model, surfaces of ribosomes contain fluid docking sites for G-quadruplex-specific proteins and foci for nucleic acid assemblies, including in polyribosomes. In addition, it has been shown that G-quadruplexes can form phase separated gels in the absence of protein components [39]. It is conceivable that polysomes are surrounded by disordered RNA gels, contributed by ribosomal tentacles and stabilized by heterogeneous G-quadruplexes.

The experiments presented in this study were performed in a cell-free environment. The extent to which rRNA G-quadruplexes form inside cells as well as their specific cellular functions remain to be determined. The 5’-UTR of NRAS, one of the most widely studied G-quadruplex-forming RNAs, contains four tandem G-tracts that have been reported to form G-quadruplexes both *in vitro* and in cells [33, 40]. We have identified ten tandem G-tracts in *tentacle a* of human ES7 rRNA and four in *tentacle b*. Our results indicate that these G-tracts form stable G-quadruplexes *in vitro*. Considering that rRNA represents over 80% of the total RNA in cells, our observation of rRNA G-quadruplexes significantly expands the possibilities for participation of G-quadruplexes in biological processes. Recent studies used G-quadruplex-RNA-specific precipitation to pull down rRNA from human cells [40].

Together, the results presented here highlight that the increase in size in rRNA expansion segments of humans has been complemented with the capability of forming G-quadruplexes. The observation of rRNA G-quadruplex regions across the phylum Chordata indicates potentially conserved functions, which most probably involve the association of specific proteins and/or assembly of polysomes.

## METHODS

ES7_HS_ spans nucleotides 436 to 1311 of the *H. sapiens* LSU rRNA and contains sequences that we call GQES7-1 (nts 583-652) and GQES7-2 (nts 825-853). RNAs corresponding to ES7_HS_, GQES7-1 and GQES7-2 were synthesized *in vitro* by transcription (HiScribe^™^T7 High Yield RNA Synthesis Kit, New England Biolabs). *mt*ES7-1 and *mt*ES7-2 were ordered as RNA oligomers. Baker’s yeast tRNAs were purchased from Roche. RNA purity was monitored by 8 M Urea 5% acrylamide gel in TBE buffer. Complete sequences of ES7_HS_, GQES7-1, GQES7-2, *mt*ES7-1 and *mt*ES7-2 are contained in Table S.1.

### HEK293T 28S rRNA extraction and purification

HEK293T cells were grown to 60% confluency after which total RNA was extracted with TRI Reagent^®^ (Sigma-Aldrich). 28S rRNA was extracted from an agarose gel by running the rRNA into wells in the center of the gel, where the rRNA was extracted with a pipette. The rRNA was precipitated in 5 M ammonium acetate-acetic Acid, pH 7.5 with excess ethanol. 28S rRNA purity was monitored on 1% agarose gels (Figure S.1).

### Thioflavin T (ThT) fluorescence

RNAs were prepared at a final concentration of 0.6 μM (strand) and annealed in 150 mM KCl, 10 mM Tris-HCl, pH 7.5, 2 μM ThT by cooling from 90°C to 25°C at 1°/min. RNAs were incubated at 4°C for 10 min and were loaded onto a Corning^®^ 384 Well Flat Clear Bottom Microplate. Fluorescence from 300-700 nm, exciting at 440 nm were acquired on a BioTek Synergy^™^ H4 Hybrid plate reader. When appropriate, pyridostatin (PDS) was added to the desired concentration after the RNA was annealed.

### Circular dichroism

RNA at 2 μM (strand) in 50 mM KCl and 10 mM Tris-HCl (pH 7.5) was annealed as described above. CD spectra were acquired at 20 °C on a Jasco J-810 spectropolarimeter using 1 mm cuvettes. Data from 200-320 nm was acquired at a rate of 100 nm/min with 1 sec response, a bandwidth of 5 nm, and averaged over three measurements. The buffer spectrum was subtracted. Smoothing was performed with Igor Pro. The observed ellipticity (θ, mdeg) was normalized [41] using the expression Δ*ε = θ*/(32,980 × *c* × *l*), where *c* is the molar strand concentration of the RNA and *l* is the path length of the cuvette in centimeters.

### Ion-dependent electrophoresis

Aliquots of RNAs at 1 μM (strand) were annealed in the presence of either Na^+^ or Li^+^ or K^+^ at various concentrations (50 mM, 100 mM, 250 mM) in 10 mM Tris-HCl, pH 7.5. Samples were mixed with glycerol (50%) and resolved on 5% native acrylamide gels, which were stained for 15 min in 0.5 μM ThT and imaged on an Azure imager c400 (Azure Biosystems).

### EMSAs

The anti-G-quadruplex BG4 antibody was purchased from Absolute Antibody (Catalog #: Ab00174-1.1). GQES7-1 (3 μM) rRNA or the negative control *mt*ES7-1 RNA were annealed in 20 mM Hepes-Tris, pH 7.5, 50 mM KCl. GQES7-1 rRNA or *mt*ES7-1 RNA were combined with various concentrations of BG4 at a final RNA concentration of 1 μM RNA (strand). RNA-protein mixtures were incubated at room temperature for 20 min in 50 mM KCl. RNA-protein interactions were analyzed by 5% native-PAGE. Gels were visualized following a dual fluorescent dye protocol [42] with a Azure imager c400 (Azure Biosystems).

### rRNA - BG4 antibody dot blotting

RNAs were annealed in the presence of 50 mM KCl and were diluted 1x, 2x and 4x. GQES7-1, GQES7-2, *mt*ES7-2, tRNA: 3.2 μM, 1.6 μM 0.8 μM. ES7_HS_: 1.4 μM, 0.7 μM, 0.35 μM. 28S rRNA: 55 nM, 27.5 nM, 13.7 nM. RNAs were loaded onto nitrocellulose membranes and dried at room temperature for 30 min. RNAs were cross-linked with the membrane in a GS Gene Linker^™^ UV Chamber (Bio Rad). The membranes were blocked for 1 h at room temperature. BG4 antibody was added (1:2,000 dilution) and incubated with gentle rocking for sixty min at room temperature. The membrane was washed for ten min twice with 1X TBST and incubated for sixty min with an appropriate fluorescent secondary antibody anti-mouse (1:10,000 dilution) (Biotium, #20065-1). The membrane was washed for ten min twice with 1X TBST and was imaged on a Li-Cor Odyssey Blot Imager. Intact 80S ribosomes and polysomes were purified from HEK293, which were incubated 5 min in 10 μg/mL cycloheximide at 37°C. Lysis buffer (10 mM NaCl, 10 mM MgCl_2_, 10 mM Tris-HCl, pH 7.5, 1% Triton X-100, 1% sodium deoxycholate, 0.2 U/mL DNase I, RNase inhibitor, 1mM dithiothreitol, 10 μg/mL cycloheximide) was used to scrap the cells. Nuclei and cell debris were removed by centrifugation and the supernatant was transferred to a 15-50% sucrose gradient containing 100 mM NaCl, 10 mM MgCl_2_, 30 mM Tris-HCl, pH 7.5 and centrifuged by ultracentrifugation. Purified 80S ribosomes and polysomes were then incubated at room temperature for 15 min in the presence of 50 mM KCl with or without 10 μM PDS. Ribosomes or polysomes were added iteratively in 30-min intervals to the same site on a nitrocellulose membrane (0.9 μg, 2.7 μg, 4.5 μg). The membrane was then treated as described above. BG4 was added to a final dilution of 1:1,000 and the secondary antibody was added to a final dilution of 1:5,000.

### Mung bean nuclease (MBN) probing

ES7_HS_ and tRNA were prepared at 100 ng/μL and annealed in the presence/absence of 100 mM KCl, 15 mM Tris-HCl (pH 7.5) by cooling from 90°C to 25°C, at 1°/min. PDS was added to the annealed RNA to a final concentration of 2 μM. One unit of MBN was added per μg of RNA and samples were incubated at 30 °C for 30 min. SDS was added to a final concentration of 0.01% to denature the nuclease and RNA was purified by ethanol precipitation. The extent of RNA cleavage was determined on an 8 M urea 5% acrylamide (19:1 acrylamide/bisacrylamide) gel stained with ethidium bromide.

### ES7 secondary structures

Secondary structures of human and *D. melanogaster* ES7 were obtained from RiboVision [43]. Nucleotides of G-quadruplex regions in *P. troglodytes, M. musculus* and *G. gallus* (Table 1) were numbered as in Bernier [1], subtracting the nucleotides from the 5.8S rRNA.

### Phylogeny and Multiple Sequence Alignments

The SEREB MSA [1] was used as a seed to align additional eukaryotic ES7 sequences to increase the density of eukaryotic species in the MSA. The 28S rRNA sequences in the SEREB MSA were used to search [44] the NCBI databases [45] for LSU rRNA sequences. The SEREB database has sequences from 10 chordate species; seven additional chordate species from 7 new orders were added to the ES7 *tentacle a* MSA (Figure 7, Table S.3). Sequences without intact ES7 *tentacle a* were excluded. Sequences with partial 28S rRNA were marked as partial. Sequences inferred from genomic scaffolds were marked as predicted (Table S.3). The extended database was queried for G-quadruplex-forming sequences.

Sequences were incorporated into the SEREB-seeded MSA using MAFFT [46] and adjusted manually using BioEdit [47]. Manual adjustments incorporated information from available secondary structures. In some cases, the positions of G-tracts in sequences with large gaps relative to *H. sapiens* are not fully determined, as they can be aligned equally well with flanking G-tracts in the MSA. Alignment visualization was done with Jalview [48]. The phylogenetic tree and the timeline of clade development were inferred from TimeTree [49].

Analysis of the entire LSU was performed on SEREB sequences, which are highly curated and always complete. This procedure ensured that negative results indicate absence of G-quadruplex-forming sequences from intact rRNAs rather than absence from rRNA fragments that lack the appropriate regions. G-quadruplex-forming sequences are not detected in any of the 20 non-chordate members of the SEREB database.

### SILAC

HEK293T cells were cultured in SILAC media - “heavy” or “light” Dulbecco’s Miodified Eagle Media (DMEM) (Thermo Scientific) supplemented with 10% dialyzed fetal bovine serum (FBS) (Corning) and 1% penicillin-streptomycin solution (Sigma) in a humidified incubator at 37 °C with 5% carbon dioxide. The heavy media contained 0.798 mM L-lysine (^13^C_6_ and ^15^N_2_, Cambridge Isotope Laboratories) and 0.398 mM L-arginine (^13^C_6_, Cambridge Isotope Laboratories). The light media had the same concentrations of normal lysine and arginine (Sigma). Media were supplemented with 0.2 mg/mL proline (Sigma) to prevent arginine-to-proline conversion. Heavy and light cells were grown for at least six generations. Once the confluency reached 80%, cells were harvested by scraping, washed twice with ice-cold PBS (Sigma), lysed in a buffer containing 10 mM HEPES pH=7.4, 200 mM potassium chloride, 1% Triton X-100, 10 mM magnesium chloride (all from Sigma) and 1 pill/10 mL cOmplete ULTRA tablet protease inhibitor (Roche), and incubated on an end-over-end shaker at 4 °C for 1 hour. Lysates were centrifuged at 25,830 g at 4 °C for 10 minutes, and the supernatants were collected and kept on ice.

Ten μg of GQES7-1-Biotin RNA was annealed as described above in the presence of 10 mM Tris-HCl, pH 7.5, and 100 mM KCl. Twenty μL of magnetic streptavidin-coated beads (GE Healthcare) were washed with Lysis buffer (10 mM HEPES, pH 7.4, 200 mM KCl, 1% Triton X-100, 10 mM MgCl_2_, protease inhibitors). Annealed RNA was then added to the washed beads and incubated at 4°C for 30 min with gentle shaking. For control experiments, no RNA was added. SILAC cell lysates were incubated with 0.5 mg *E. coli* tRNA per 1 mg protein at 4°C for 30 min with gentle shaking. “RNA+Beads” and control “Beads Only” samples were transferred into the SILAC cell lysates: “RNA+Beads” were added to the Heavy cell lysate and “Beads Only” was added to “Light” HEK293T cell lysate. As a replicate, “RNA+Beads” was added “Light” cell lysate and “Beads Only” was added to “Heavy” cell lysate. 200 U/mL of RNasin was added and the lysates were incubated at 4 °C for 2 hrs with gentle shaking. Samples were centrifuged, the supernatant was discarded, and the pelleted beads were washed with lysis buffer with increasing KCl concentrations (0.4 M, 0.8 M, 1.6 M). After the three washes, 100 μL of the elution buffer (100 mM Tris-HCl, pH 7.4,1% SDS, 100 mM DTT) was added to one of the two samples and then combined with the beads of the corresponding sample. “RNA+Beads” in “Heavy” lysates were combined with “Beads Only” in “Light” lysates and “RNA+Beads” in “Light” lysates were combined with “Beads Only” in “Heavy” lysates. The combined samples were boiled and then briefly centrifuged. Beads were discarded and samples were analyzed with an online LC-MS system.

### Mass spectrometry analyses

Eluted proteins were diluted 10 times with 50 mM HEPES pH=7.4 and were alkylated with 28 mM iodoacetamide (Sigma) for 30 minutes at room temperature in the dark. Proteins were precipitated by methanol-chloroform, and the pellets were resuspended in digestion buffer containing 50 mM HEPES pH=8.8, 1.6 M urea, and 5% acetonitrile (ACN) (all from Sigma). After digestion with sequencing-grade modified trypsin (Promega) at 37 °C for 16 hours, reactions were quenched with 1% trifluoroacetic acid (TFA, Fisher Scientific) and purified with StageTip. Peptides were dissolved in 10 μL 5% ACN and 4% FA solution, and 1 μL was loaded to a Dionex UltiMate 3000 UHPLC system (Thermo Fisher Scientific) with a microcapillary column packed inhouse with C18 beads (Magic C18AQ, 5 mm, 200 Ǻ 100 mm 16 cm). A 110-minute gradient of 3-22% ACN containing 0.125% FA was used. The peptides were detected with an LTQ Orbitrap Elite Hybrid Mass Spectrometer (Thermo Fisher Scientific) controlled by Xcalibur software (version 3.0.63). MS/MS analysis was performed with a data-dependent Top20 method. For each cycle, a full MS scan in the Orbitrap with the automatic gain control (AGC) target of 10^6^ and the resolution of 60,000 at 400 m/z was followed by up to 20 MS/MS scans in the LTQ for the most intense ions. Selected ions were excluded from further sequencing for 90 seconds. Ions with singly or unassigned charge were not sequenced. Maximum ion accumulation times were 1,000 ms for each full MS scan and 50 ms for each MS/MS scans. The spectra were searched against a human protein database downloaded from UniProt using the SEQUEST algorithm (version 28) [50]. The following parameters were used: 20 ppm precursor mass tolerance; 1.0 Da fragment ion mass tolerance; trypsin digestion; maximum of 3 missed cleavages; differential modifications for methionine oxidation (+15.9949 Da), heavy lysine (+8.0142 Da), and heavy arginine (+6.0201 Da); fixed modification for cysteine carbamidomethylation (+57.0215 Da). The false discovery rates (FDR) were evaluated and controlled by the target-decoy method. Linear discriminant analysis (LDA) was used to filtered the peptides to <1% FDR based on parameters such as XCorr, differential sequence ΔC_n_, and precursor mass error. An additional filter was used to control the protein FDR to <1%. For SILAC quantification, the S/N ratios of both heavy and light peptides must be greater than 3. Otherwise, one of the two versions of the peptides (heavy or light) must have the S/N ratio greater than 10. Other peptides that did not pass these criteria were removed. The final protein ratio was calculated from the median value of the peptides from each parent protein. The raw files are publicly accessible at http://www.peptideatlas.org/PASS/PASS01260, Username: PASS01260, Password: TL3854zn.

### Western Blotting

Samples were electrophoresed on 12% SDS-PAGE gels and transferred to a nitrocellulose membrane overnight. Membranes were blocked for 1 hour at room temperature with gentle shaking and then incubated for another hour with primary antibodies: 1:200 dilution of FIP1 (mouse monoclonal, sc-398392), DDX3 (mouse monoclonal, sc-81247), FUS (mouse monoclonal, sc-47711), or hnRNP H (mouse monoclonal, sc-32310). Antibodies were obtained from Santa Cruz Biotechnology. Membranes were washed three times with 1X TBST and secondary antibody CF680 goat anti-mouse IgG (H+L) (Biotium, 20065) was added (1:5,000 dilution). Membranes were washed three times with 1X TBST and imaged on a Li-Cor Odyssey Blot Imager.

## ACKNOWLEDGEMENTS

The authors thank Dr. Jonathan B. Chaires for helpful discussions and Dr. Lizzette M. Gómez Ramos for designing the negative G-quadruplex controls. Purified 80S ribosomes and polysomes were a gift from Immagina BioTechnology. This work was supported by NASA (NNX16AJ28G and NNX16AJ29G to LDW) and the National Institutes of Health (R01GM118803 to RW).

## CONFLICT OF INTEREST

The authors declare that they have no conflict of interest with the contents of this article.

## AUTHOR CONTRIBUTIONS

SMF, SS, CI, ASP, RMW, RW, and LDW conceived and designed the experiments; SMF, SS and CI performed the experiments; ASP and PIP conducted the phylogenetic analysis. SMF, PIP, SS, ASP, RMW, RW, and LDW analyzed data; SMF, PIP, SS and LDW prepared figures; and SMF, PIP and LDW wrote the paper.

